# The therapeutic effects of intratumoral injection of IL12IL2GMCSF fusion protein on canine tumors

**DOI:** 10.1101/2020.06.03.131904

**Authors:** Xiaobo Du, Bin Zhang, Fan Hu, Chuantao Xie, Haiyan Tian, Qiang Gu, Haohan Gong, Xiaoguang Bai, Jinyu Zhang

**Author notes:** To whom correspondence should be addressed: Jinyu Zhang, Beijing Kenuokefu Biotechnology Company, Building 1, 29#, Shengmingyuan Road, Changping district, Beijing, 102200, China.

## Abstract

Regulating the immune system through tumor immunotherapy to defeat tumors is currently one of the most popular methods of tumor treatment. We previously found that the combinations of IL12, IL2 and GMCSF has superior antitumor activities. In this study, IL12IL2GMCSF fusion protein was produced from 293 cells transduced by expression lentiviral vector. IL12IL2GMCSF fusion protein was injected into canine tumors, and the efficacy was monitored to evaluate the potential of tumor immunotherapy. In 14 canines injected with IL12IL2GMCSF protein, 6 had a complete response (CR), 7 had a partial response (PR), and 1 had stable disease (SD). IL12IL2GMCSF fusion protein has great anti-tumor potential. Intratumoral injection of IL12IL2GMCSF can effectively treat tumors without severe adverse effects. It provides a promising immune therapeutics in canine cancers.

## 1 Introduction

The occurrence and development of tumors are complex and dynamic processes. With further research, researchers have begun to realize that in addition to the tumor cells, the physiologic state of the tumor microenvironment, including the surrounding stromal cells, extracellular matrix, and signal molecules, is closely related to tumor occurrence and development (1). The importance of the tumor microenvironment in the genesis and development of tumors suggests that targeting the tumor microenvironment may be one of the most effective strategies to eradicate cancer cells.

Cytokines are signaling proteins that have important effects on cell-to-cell interactions and communications (2). Cytokines are released under various cellular stressors, such as infection, inflammation, and cell damage. Studies have shown that cytokines produced in the tumor microenvironment are closely related to the occurrence and development of tumors (3–6). Interleukin(IL) 2 is an important cytokine with multiple effects on the immune system. As an effective activator of cytotoxic T cells and NK cells, IL2 is considered to have anti-tumor effects and has been approved by the FDA as a tumor immunotherapy with specific anti-tumor effects (7–9). Like IL2, IL12 can also activate T cells and NK cells. IL12 promotes the differentiation of CD4 + T cells into Th1 CD4 + T cells and enhances the activity of CD8 + cytotoxic T lymphocytes (10–12). IL12 has also been designated as one of the ideal candidates for tumor immunotherapy and has shown tumor suppressive effects in melanoma, colon cancer, and breast cancer models (13–15). Granulocyte macrophage-colony stimulating factor (GMCSF) belongs to the hematopoietic cytokine family and can promote the recruitment and activation of bone marrow cells (16). Clinical studies have shown that GMCSF is an immunopotentiator and is a very promising anti-tumor drug (17).

Cytokines, such as IL12, IL2, and GMCSF, not only show anti-tumor activity when used alone, but also show stronger anti-tumor activity when used in combination (18–20); however, the antitumor effect of cytokines is affected by multiple factors, such as dosage, method of administration, time, and medication strategy. In this study we evaluated the therapeutic effects of intratumoral injection of IL12, IL2, and GMCSF fusion proteins on malignant tumors in canines to provide experimental evidence for the discovery and development of tumor immunotherapy drugs.

## 2 Material and Methods

### 2.1 Study design

This study was conducted simultaneously in several animal hospitals in Beijing from October 2017 to December 2019. We obtained approvals from Ethics Committees of each animal hospital and consents from all dog owners. All dogs selected were out of other treatment options and their treatments could be discontinued if their owners wish to withdraw. The treatment cost was borne by the researchers. Inclusion criteria: primary lesions were visible or touchable and can be injected intratumorally. Tumors were diagnosed by histopathology. Tumor staging standards follow the WHO guidelines.

### 2.2 Production of IL12IL2GMCSF fusion protein

The IL12IL2GMCSF fusion protein consists of mature canine IL12b, IL12a, IL2 and GMCSF sequences and these subunits were tandemly connected with (G4S)3 linkers. The DNA sequence encoding this protein was synthesized and subcloned into vector pLentis-CMV-MCS-IRES-PURO between BamHI and XhoI sites, generating expression vectors pLentis-CMV-IL12IL2GMCSF-IRES-PURO. A secretory signal peptide was placed upstream of the fusion protein, and a 6*his peptide was placed downstream for affinity purification. The lentiviral particles were produced by cotransfection of pMD2.G, psPAX2 and pLentis-CMV-IL12IL2GMCSF-IRES-PURO into 293FT cells. 293 cells were transduced with pLentis-CMV-IL12IL2GMCSF-IRES-PURO lentivirus and selected with 3 ug/ml puromycin, generating cells line 293(IL12IL2GMCSF) for stable expression of IL12IL2GMCSF protein. The protein was collected from the supernatant of cultured cells and concentrated by Amicon Ultra-15 Centrifugal Filter Unit (Merck). Then the fusion protein was purified using BeaverBeads IDA-Nickel Kit (Beaverbio), according to the manufacturer’s instructions. The concentration of the fusion protein was measured with canine IL12p40 ELISA kit (R&D Systems). The purity was evaluated by SDS page. The purified protein was diluted to 500 ug/ml and stored at −20 °C.

### 2.3 Treatment of dogs with cancer

The dosage was determined by the measured tumor area. When the area is less than or equal to 6cm^2^, a single injection of 250μg was used. When the area is between 6 and 15cm^2^, a single injection of 500μg was used. When the area is greater than 15cm2, a single injection of 750μg was used. Before injection, each 0.5ml of 500μg / ml fusion protein solution was mixed well with an equal volume of glycerin to prepare an injection solution of about 1ml. The subjects received injection once every two weeks and the injection volume was adjusted based on the tumor area. After the injection, animals stay in the hospital for half an hour and were taken home by their owners to observe body temperature, mental state, appetite, urine and other conditions.

### 2.4 Assessment of therapeutic effects and side effects

The size of the tumor is measured with a caliper. Generally, it is measured once a week, and some difficult to measurement sites (such as cases that can only be measured after anesthesia) are measured once every two weeks or longer. How to measure the tumor diameter and how often to measure are determined by the response changes in tumor diameter. Response standards follow Standard Response Evaluation Criteria in Solid Tumors (RECIST). Disappearance of the lesion is a complete response (CR), a reduction in tumor diameter of more than 30% is a partial response (PR), and a change in tumor diameter between 30% decrease and 20% increase is considered stable disease (SD).

The evaluation of adverse events follows the Veterinary Cooperative Oncology Group-Common Terminology Criteria for Adverse Events (VCOG-CTCAE) v1.1 guidelines. It was carried out through hospital examination and observation by the owners, including the mental state of the dogs, eating and defecation, body temperature, tumor appearance and blood biochemical indicators.

### 2.5 Statistical analysis

The Kaplan-Meier method was used to calculate overall survival (OS) curves. All analysis was performed using GraphPad Prism 5 software (GraphPad Software, Inc., LaJolla, CA, USA).

## 3 Results

### 3.1 Cases

Fourteen dogs, 9-15 years old (age unknown for canine #11) with malignant tumors, including sarcomas, breast cancer, melanomas, thyroid cancer, mast cell cancer, fibrosarcomas, transmissible venereal tumor (TVTs), and perianal tumors, were included in this study. The canine characteristics, such as breed, age, tumor type, treatment history, and tumor stage before using IL12IL2GMCSF fusion protein treatment, are summarized in Table 1.

**Table 1.** Clinical characteristics of canine tumor cases treated with IL12IL2GMCSF fusion proteins.

| Canine No. | Breed | gender | weight (kg) | Tumor type | Age (Year) | Tumor size (cm<sup>2</sup>) | Tumor character | Stage |
| --- | --- | --- | --- | --- | --- | --- | --- | --- |
| 1 | Mix | M | 5 | Sarcoma | 9 | 7 | Primary tumor | III |
| 2 | Beagle | F | 12 | Breast Cancer | 11 | 3 | Recurrence after surgical resection | IV |
| 3 | Mixed | M | 10 | Sarcoma | 10 | 6 | Primary tumor | II |
| 4 | Mixed | F | 15 | Breast cancer | 15 | 4 | Recurrence after surgical resection | IV |
| 5 | Chihuahua | SF | 3 | Hemagiopericytoma | 11 | 42 | Primary tumor | III |
| 6 | Mixed | M | 10 | Melanoma | 13 | 1 | Recurrence after surgical resection | I |
| 7 | Samoyed | F | 22 | Thyroid cancer | 11 | 126 | Primary tumor | III |
| 8 | Pekingese | CM | 5 | Mast cell tumor | 14 | 15 | Three surgeries before treatment | IV |
| 9 | Pomeranian | F | 2.5 | TVT | 15 | 6 | Primary tumor | III |
| 10 | Afghan hound | SF | 27 | Mast cell tumor | 13 | 30 | Primary tumor | III |
| 11 | Beagle | F | 11 | Fibrosarcoma | 12 | 24 | Primary tumor | II |
| 12 | Alaskan Malamute | M | 50 | Perianal tumor | 12 | 10 | Primary tumor | II |
| 13 | Chow Chow | M | 32 | Perianal tumor | 11 | 8 | Primary tumor | II |
| 14 | Sheltie | SF | 21 | Fibrosarcoma | 12 | 15 | Recurrence after surgical resection | III |

### 3.2 Production of IL12IL2GMCSF fusion protein

The schematic structure of IL12IL2GMCSF fusion protein was shown in Figure 1a. The concentration in culture supernatant was 106.5 ng/ul and it was 1265.3 ng/ul after purification (Figure 1b). The purity of monomer protein is ~ 90.3% (Figure 1c).

**Figure 1.**
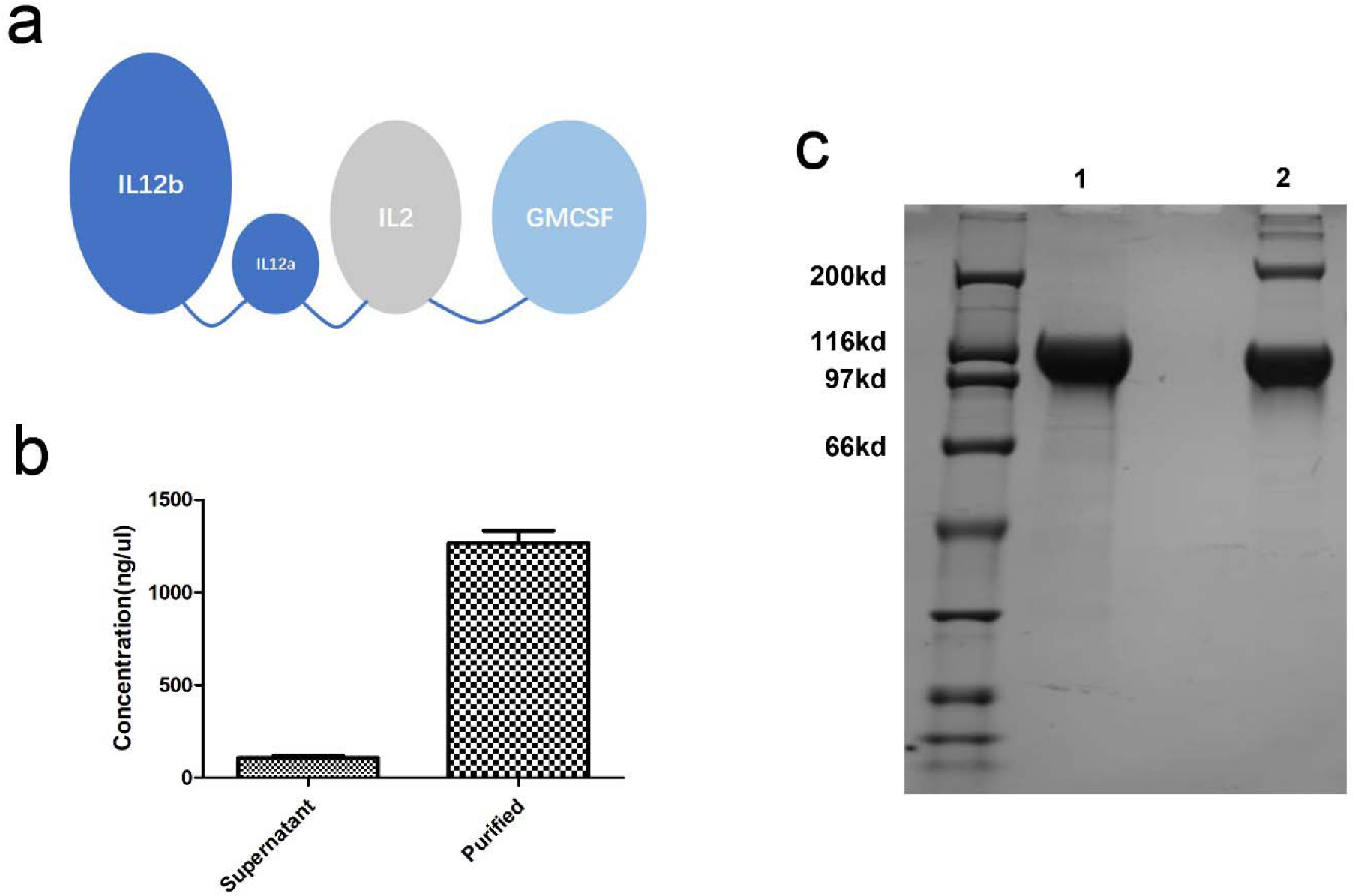
Production of IL12IL2GMCSF fusion protein. (a) The schematic structure of IL12IL2GMCSF fusion protein. (b) Detection of IL12IL2GMCSF fusion protein level by ELISA. (c) Gel electrophoresis in purified IL12IL2GMCSF fusion protein in SDS PAGE. 1 Reduced condition; 2 Non-reduced condition.

### 3.3 Treatment of dogs

Table 2 showed the tumor sites in dogs receiving intratumoral injections of IL12IL2GMCSF fusion proteins. Dosing cycles and doses was shown in Table 2. The efficacy of intratumoral injections of IL12IL2GMCSF fusion proteins was shown in Table 2. Six canines had a complete response (CR), seven had a partial response (PR) and one had stable disease (SD).

**Table 2.**
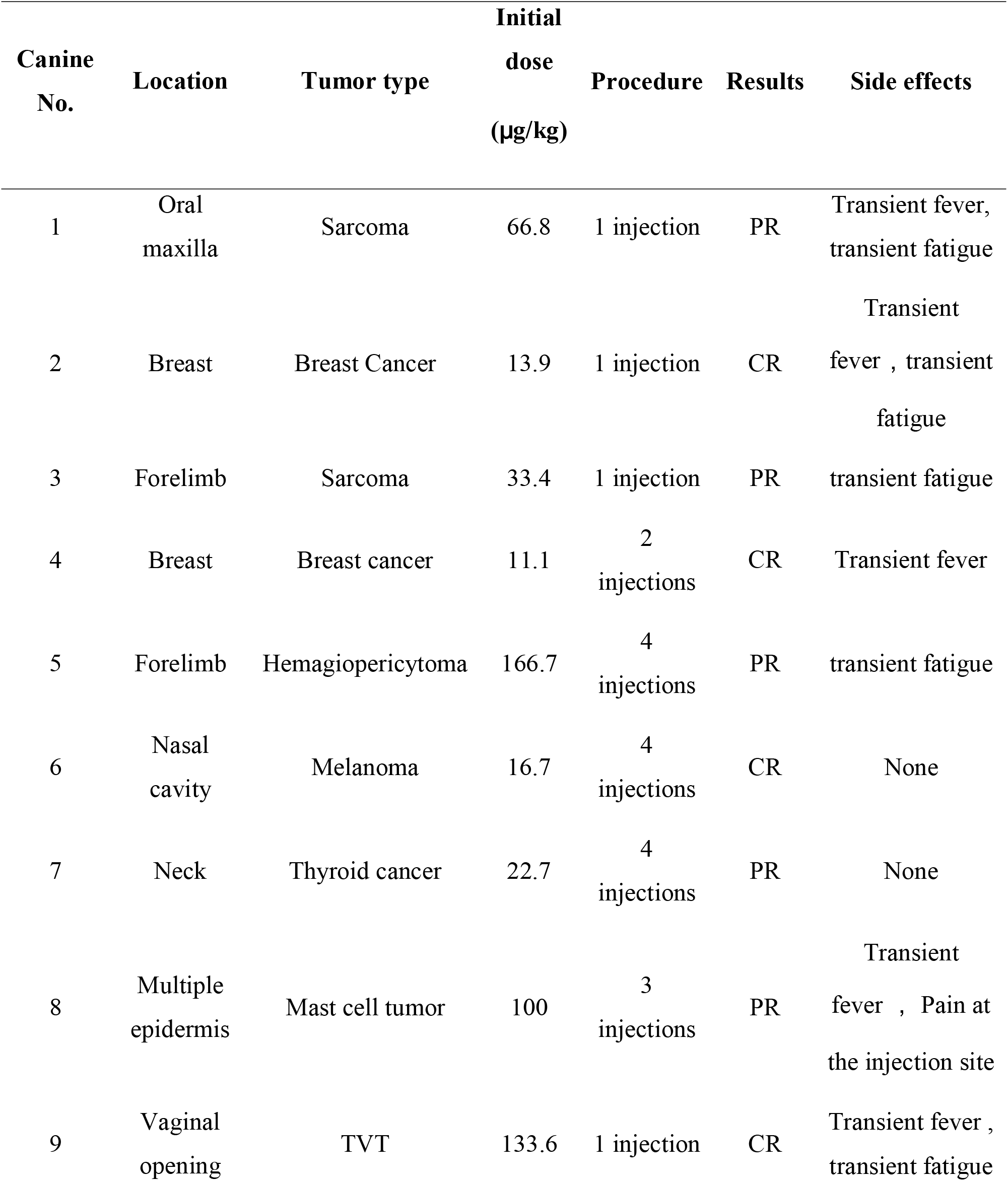
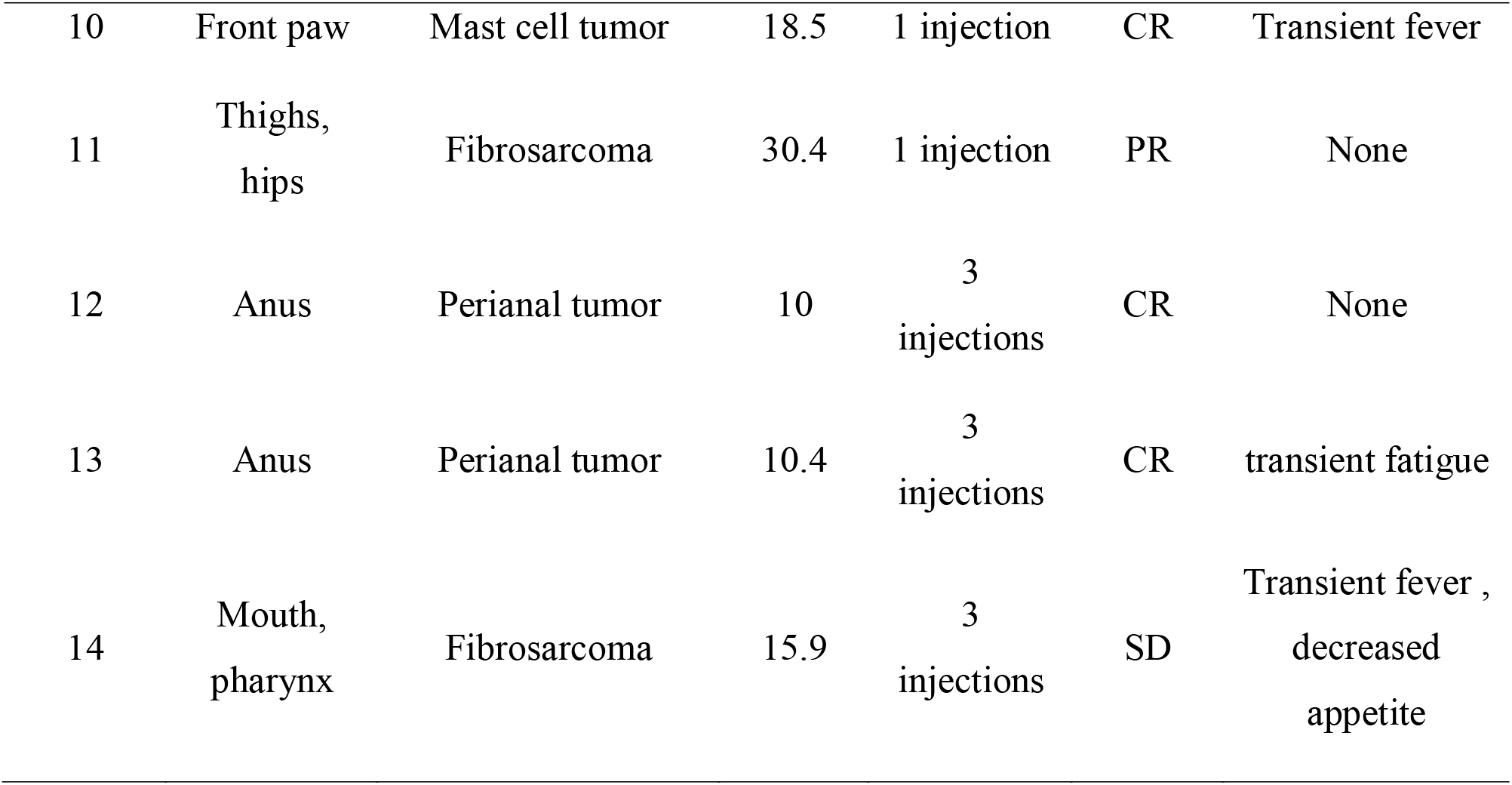
Effects of intratumoral injection of IL12IL2GMCSF fusion protein to treat canine tumors.

### 3.4 Clinical outcomes

Of the 12 canines that were followed, the shortest survival time after administration of IL12IL2GMCSF was 178 days. Tumor regression in the dog with melanoma or leg tumor after treatment was shown in Figure 2 and 3. The Kaplan-Meier survival curve is shown in Figure 4. Upon completion of the study, 5 of 12 canines we were able to track had died (canine nos. 2, 3, 7, 8, and 10) and 7 survived (canine nos. 4, 5, 6, 11, 12, 13, and 14). Of the 5 deaths, 2 (canine nos. 2 and 8) died of unknown causes, 1 (canine nos. 7) died of diabetes, 1 (canine nos. 3) was euthanized, and 1 (canine no. 10) died of a mass in the lung. Because canine mast cell tumors were rare in metastasizing to the lungs, the likelihood of recurrence and metastasis after treatment is low.

**Figure 2.**
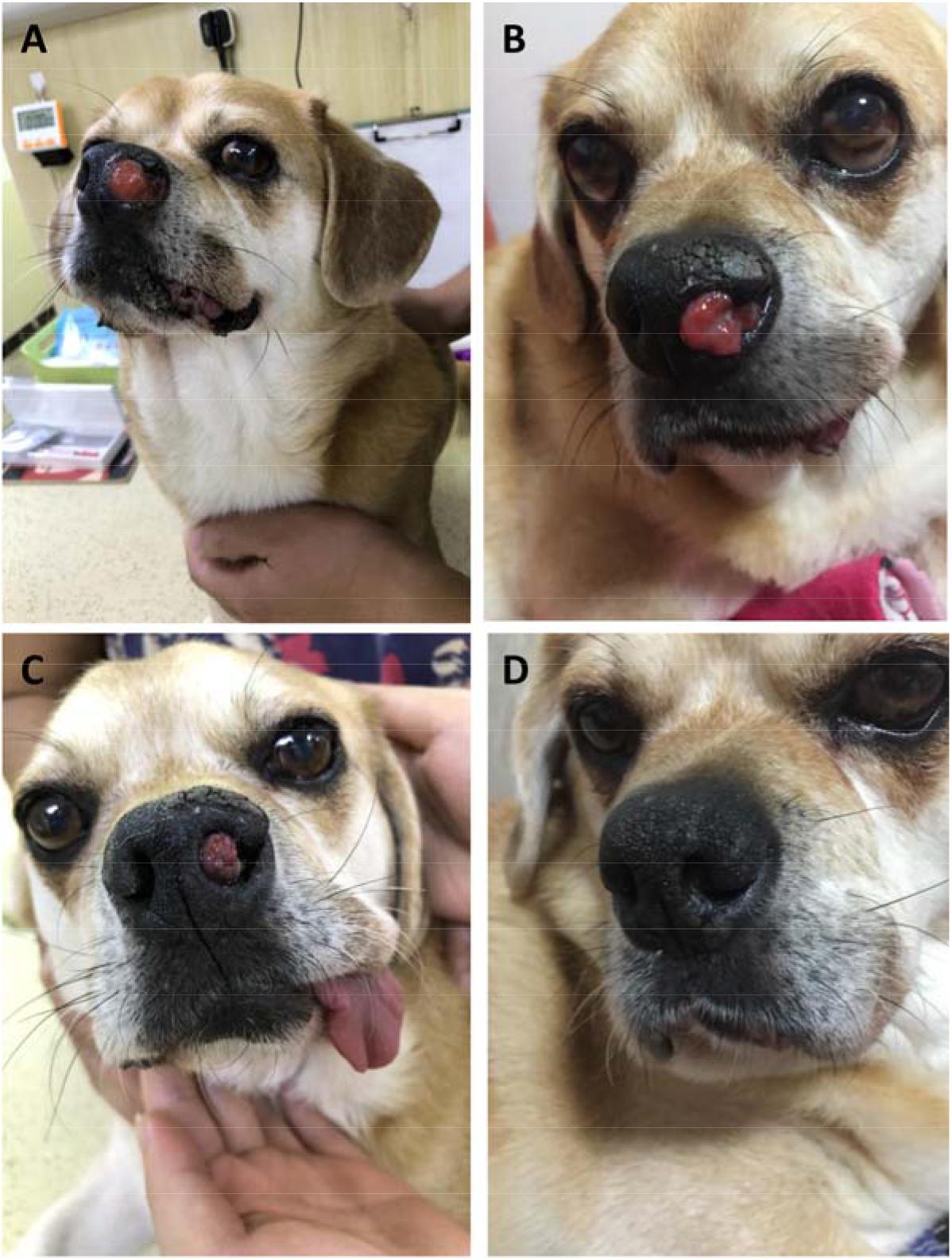
Photographs of the pre-treatment melanoma and the changes in tumors on days 30,54,360 post-treatment in case 6. A, pre-treatment (Day0)); B, Day 30; C, Day54; D, Day 360;

**Figure 3.**
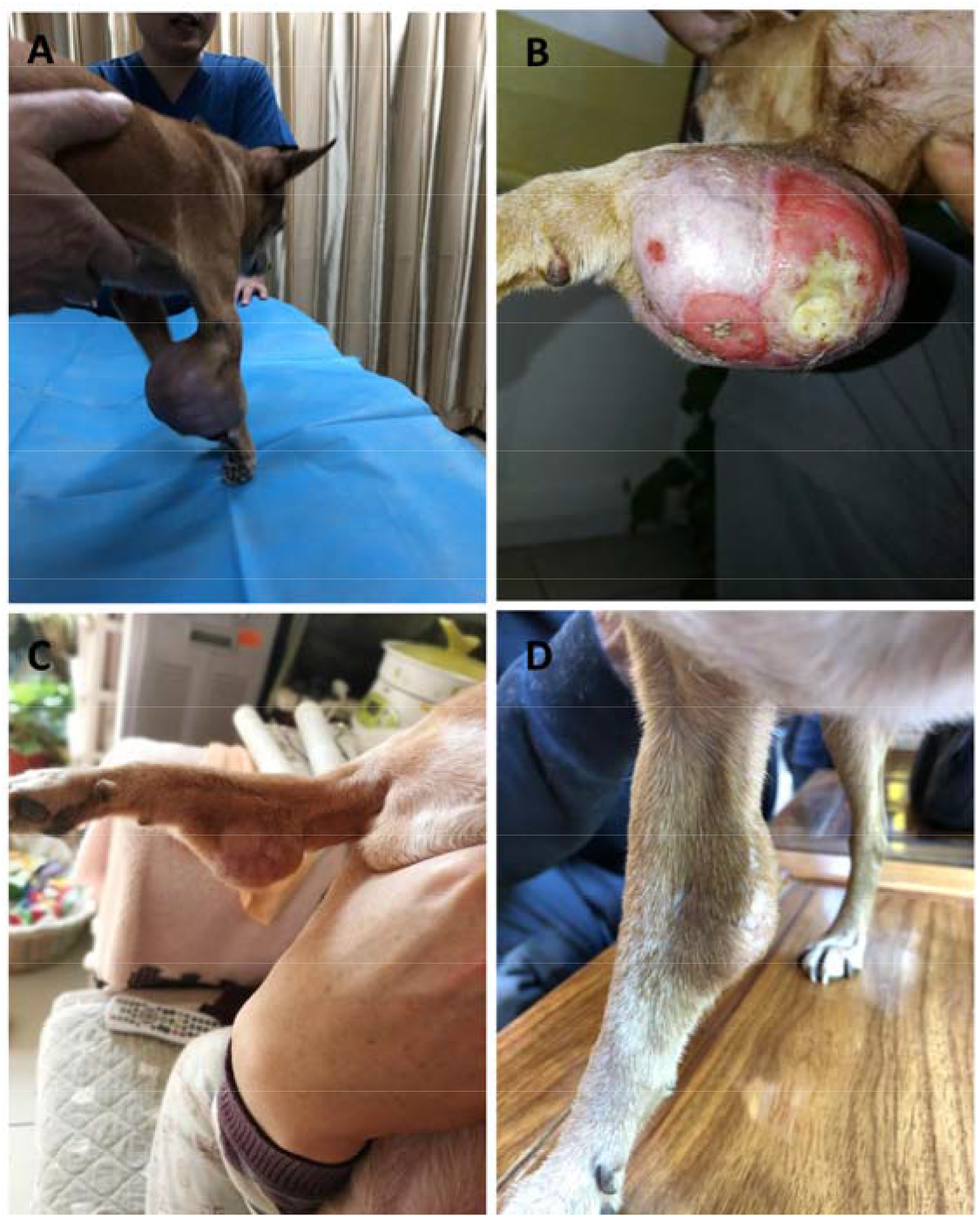
Photographs of the pre-treatment leg tumor and the changes in tumors on days 7,65,330 post-treatment in case (No. 5). A, pre-treatment (Day0)); B, Day 7; C, Day65; D, Day 330

**Figure 4.**
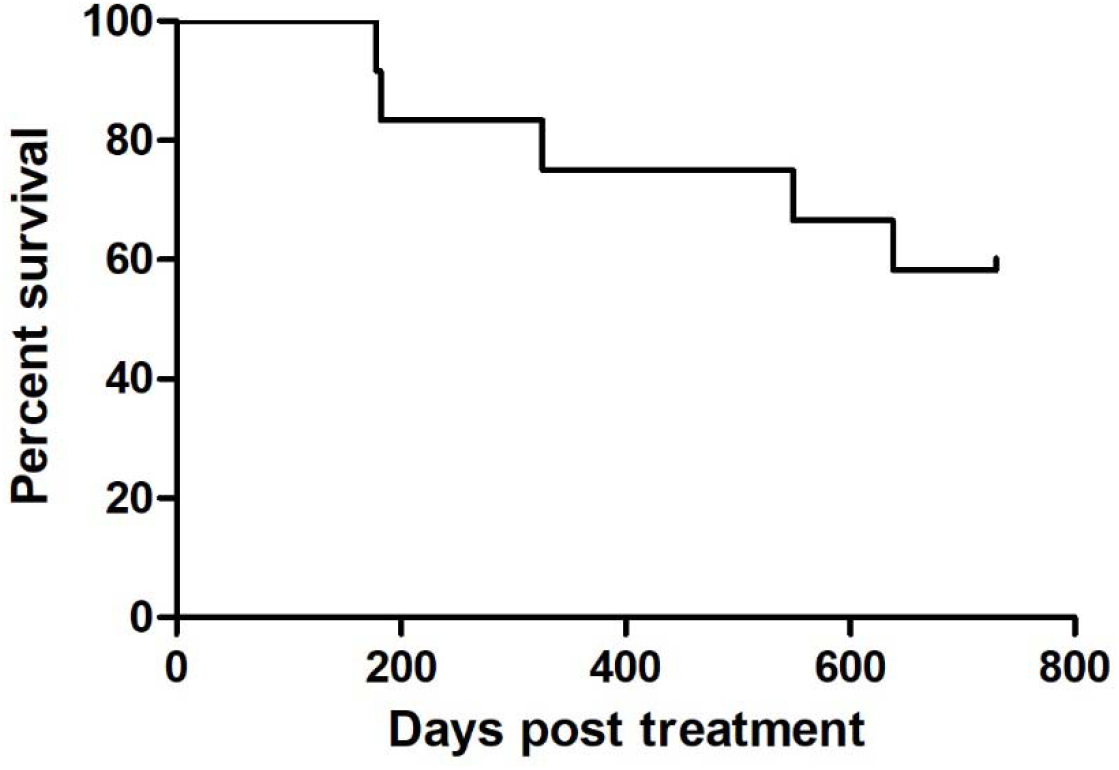
Kaplan-Meier curve depicting overall survival of dogs treated with IL12IL2GMCSF fusion protein (median 566 days, n = 12).

### 3.5 Adverse events of IL12IL2GMCSF fusion protein injection

As shown in Table 2, after the injection of IL12IL2GMCSF fusion protein,7 canines (canine nos. 1, 2, 4, 8, 9, 10 and 14) developed transient fevers, 5 canines (canine nos. 1, 2, 3, 5 and 9) developed transient fatigue, the dog no.14 showed decreased appetite and 1 canine (canine no. 8) had pain at the injection site. None of the dogs showed serious changes in blood or biochemical indicators. No adverse reactions requiring symptomatic treatment were observed.

## 4 Discussion

The theory that combined application of different cytokines in the tumor environment can lead to a strong synergistic anti-tumor response provides an effective strategy for immunotherapy of malignant tumors (19, 21). In this study, intratumoral injections of IL12IL2GMCSF fusion proteins were used to evaluate the therapeutic effect of the combined application of the above cytokines on canine tumors. It was found that IL12IL2GMCSF fusion protein exhibited various degrees of anti-tumor immunity in a variety of malignant tumors, such as sarcomas, breast cancer, perianal tumors, fibrosarcomas, mast cell tumors, melanomas, and TVTs. There were often large differences between different types of tumors, especially with respect to genomic mutations and immune cell infiltration (22). The effectiveness of IL12IL2GMCSF fusion proteins in a variety of malignancies indicated that IL12IL2GMCSF fusion proteins had a wide range of anti-tumor immune activities and were a potentially effective tumor immunotherapy.

Strategies to regulate the immune system have made great progress in the field of anti-tumor therapy, but drugs that activate the immune system are also likely to cause severe non-specific systemic inflammation and autoimmune side effects (23). Clinical studies have shown that high doses of IL2 for metastatic melanoma are likely to cause dose-dependent cytotoxicity (24). Injecting immunomodulatory drugs directly into accessible lesions (primary or metastatic tumors) is a simple way to reduce immunotoxicity. The rationale for this method is that local administration will preferentially remain at the site of the injected tumor, and this retention may be advantageous if local administration allows administration at lower doses than systemic administration (23). Local administration has been cited in the treatment of various tumors, such as melanomas, breast cancer, ovarian cancer, lymphomas, and bladder cancer, and has effectively reduced adverse reactions (25–30). Based on the results of previous studies, we also used intratumoral injections. The results of this study showed that among the dogs that received intratumoral injections of IL12IL2GMCSF fusion proteins at the dose used in this study, there were no serious adverse reactions except for four dogs (canine nos. 1, 2, 8, 10) who developed transient fever and one (case 8) also had injection site pain.

In summary, the current research results showed that IL12IL2GMCSF fusion protein has extensive anti-tumor immunotherapy potential. Intratumoral injection of the fusion protein can effectively treat tumors without severe adverse reaction. This study has provided a powerful experimental foundation for the discovery and development of new drugs for tumor immunotherapy.

## References

1. Wang M, Zhao J, Zhang L, Wei F, Lian Y, Wu Y, Gong Z, Zhang S, Zhou J, Cao K, et al. Role of tumor microenvironment in tumorigenesis. Journal of Cancer. 2017;8(5):761–73.

2. Zhang JM, and An J. Cytokines, inflammation, and pain. International anesthesiology clinics. 2007;45(2):27–37.

3. Mantovani A, Barajon I, and Garlanda C. IL-1 and IL-1 regulatory pathways in cancer progression and therapy. Immunological reviews. 2018;281(1):57–61.

4. Dranoff G, Jaffee E, Lazenby A, Golumbek P, Levitsky H, Brose K, Jackson V, Hamada H, Pardoll D, and Mulligan RC. Vaccination with irradiated tumor cells engineered to secrete murine granulocyte-macrophage colony-stimulating factor stimulates potent, specific, and long-lasting anti-tumor immunity. Proceedings of the National Academy of Sciences of the United States of America. 1993;90(8):3539–43.

5. Floros T, and Tarhini AA. Anticancer Cytokines: Biology and Clinical Effects of Interferon-alpha2, Interleukin (IL)-2, IL-15, IL-21, and IL-12. Seminars in oncology. 2015;42(4):539–48.

6. Lee S, and Margolin K. Cytokines in cancer immunotherapy. Cancers. 2011;3(4):3856–93.

7. Rosenberg SA, Lotze MT, Muul LM, Leitman S, Chang AE, Ettinghausen SE, Matory YL, Skibber JM, Shiloni E, Vetto JT, et al. Observations on the systemic administration of autologous lymphokine-activated killer cells and recombinant interleukin-2 to patients with metastatic cancer. The New England journal of medicine. 1985;313(23):1485–92.

8. Jiang T, Zhou C, and Ren S. Role of IL-2 in cancer immunotherapy. Oncoimmunology. 2016;5(6):e1163462.

9. Sun Z, and Ren Z. A next-generation tumor-targeting IL-2 preferentially promotes tumorinfiltrating CD8(+) T-cell response and effective tumor control. 2019;10(1):3874.

10. Chehimi J, Starr SE, Frank I, Rengaraju M, Jackson SJ, Llanes C, Kobayashi M, Perussia B, Young D, Nickbarg E, et al. Natural killer (NK) cell stimulatory factor increases the cytotoxic activity of NK cells from both healthy donors and human immunodeficiency virus-infected patients. The Journal of experimental medicine. 1992;175(3):789–96.

11. Perussia B, Chan SH, D’Andrea A, Tsuji K, Santoli D, Pospisil M, Young D, Wolf SF, and Trinchieri G. Natural killer (NK) cell stimulatory factor or IL-12 has differential effects on the proliferation of TCR-alpha beta+, TCR-gamma delta+ T lymphocytes, and NK cells. Journal of immunology (Baltimore, Md: 1950). 1992;149(11):3495–502.

12. Chan SH, Kobayashi M, Santoli D, Perussia B, and Trinchieri G. Mechanisms of IFN-gamma induction by natural killer cell stimulatory factor (NKSF/IL-12). Role of transcription and mRNA stability in the synergistic interaction between NKSF and IL-2. Journal of immunology (Baltimore, Md: 1950). 1992;148(1):92–8.

13. Hwang MP, Fecek RJ, Qin T, Storkus WJ, and Wang Y. Single injection of IL-12 coacervate as an effective therapy against B16-F10 melanoma in mice. Journal of controlled release: official journal of the Controlled Release Society. 2019;318(270–8.

14. Men K, Huang R, Zhang X, Zhang R, Zhang Y, He M, Tong R, Yang L, Wei Y, and Duan X. Local and Systemic Delivery of Interleukin-12 Gene by Cationic Micelles for Cancer Immunogene Therapy. Journal of biomedical nanotechnology. 2018;14(10):1719–30.

15. Yang SX, Wei WS, Ouyan QW, Jiang QH, Zou YF, Qu W, Tu JH, Zhou ZB, Ding HL, Xie CW, et al. Interleukin-12 activated CD8(+) T cells induces apoptosis in breast cancer cells and reduces tumor growth. Biomedicine & pharmacotherapy = Biomedecine & pharmacotherapie. 2016;84(1466–71.

16. Ushach I, and Zlotnik A. Biological role of granulocyte macrophage colony-stimulating factor (GM-CSF) and macrophage colony-stimulating factor (M-CSF) on cells of the myeloid lineage. Journal of leukocyte biology. 2016;100(3):481–9.

17. Dranoff G. GM-CSF-secreting melanoma vaccines. Oncogene. 2003;22(20):3188–92.

18. Wen Q, Xiong W, He J, Zhang S, Du X, Liu S, Wang J, Zhou M, and Ma L. Fusion cytokine IL-2-GMCSF enhances anticancer immune responses through promoting cellcell interactions. Journal of translational medicine. 2016;14(41.

19. Zhang J, Jiang H, and Zhang H. In situ administration of cytokine combinations induces tumor regression in mice. EBioMedicine. 2018;37(38–46.

20. Gollob JA, Veenstra KG, Parker RA, Mier JW, McDermott DF, Clancy D, Tutin L, Koon H, and Atkins MB. Phase I trial of concurrent twice-weekly recombinant human interleukin-12 plus low-dose IL-2 in patients with melanoma or renal cell carcinoma. Journal of clinical oncology: official journal of the American Society of Clinical Oncology. 2003;21(13):2564–73.

21. Weiss JM, Subleski JJ, Wigginton JM, and Wiltrout RH. Immunotherapy of cancer by IL-12-based cytokine combinations. Expert opinion on biological therapy. 2007;7(11):1705–21.

22. Meacham CE, and Morrison SJ. Tumour heterogeneity and cancer cell plasticity. Nature. 2013;501(7467):328–37.

23. Milling L, Zhang Y, and Irvine DJ. Delivering safer immunotherapies for cancer. Advanced drug delivery reviews. 2017;114(79–101.

24. Alwan LM, Grossmann K, Sageser D, Van Atta J, Agarwal N, and Gilreath JA. Comparison of acute toxicity and mortality after two different dosing regimens of high-dose interleukin-2 for patients with metastatic melanoma. Targeted oncology. 2014;9(1):63–71.

25. Stearns V, Mori T, Jacobs LK, Khouri NF, Gabrielson E, Yoshida T, Kominsky SL, Huso DL, Jeter S, Powers P, et al. Preclinical and clinical evaluation of intraductally administered agents in early breast cancer. Science translational medicine. 2011;3(106):106ra8.

26. Galanis E, Hartmann LC, Cliby WA, Long HJ, Peethambaram PP, Barrette BA, Kaur JS, Haluska PJ, Jr., Aderca I, Zollman PJ, et al. Phase I trial of intraperitoneal administration of an oncolytic measles virus strain engineered to express carcinoembryonic antigen for recurrent ovarian cancer. Cancer research. 2010;70(3):875–82.

27. Brody JD, Ai WZ, Czerwinski DK, Torchia JA, Levy M, Advani RH, Kim YH, Hoppe RT, Knox SJ, Shin LK, et al. In situ vaccination with a TLR9 agonist induces systemic lymphoma regression: a phase I/II study. Journal of clinical oncology: official journal of the American Society of Clinical Oncology. 2010;28(28):4324–32.

28. Biot C, Rentsch CA, Gsponer JR, Birkhauser FD, Jusforgues-Saklani H, Lemaitre F, Auriau C, Bachmann A, Bousso P, Demangel C, et al. Preexisting BCG-specific T cells improve intravesical immunotherapy for bladder cancer. Science translational medicine. 2012;4(137):137ra72.

29. Posch C, Weihsengruber F, Bartsch K, Feichtenschlager V, Sanlorenzo M, Vujic I, Monshi B, Ortiz-Urda S, and Rappersberger K. Low-dose inhalation of interleukin-2 biochemotherapy for the treatment of pulmonary metastases in melanoma patients. British journal of cancer. 2014;110(6):1427–32.

30. Weide B, Eigentler TK, Pflugfelder A, Zelba H, Martens A, Pawelec G, Giovannoni L, Ruffini PA, Elia G, Neri D, et al. Intralesional treatment of stage III metastatic melanoma patients with L19-IL2 results in sustained clinical and systemic immunologic responses. Cancer immunology research. 2014; 2(7): 668–78.

